# The functional organization of high-level visual cortex determines the representation of complex visual stimuli

**DOI:** 10.1101/2019.12.22.852293

**Authors:** Libi Kliger, Galit Yovel

## Abstract

A hallmark of high-level visual cortex is its functional organization of neighboring clusters of neurons that are selective to single categories such as faces, bodies and objects. However, visual scenes are typically composed of multiple categories. How does category-selective cortex represent such complex stimuli? According to a normalization mechanism, the response of a single neuron to multiple stimuli is normalized by the response of its neighboring neurons (normalization pool). Here we show that category-selectivity, measured with fMRI, can provide an estimate for the heterogeneity of the normalization pool, which determines the response to multiple stimuli. These results provide a general framework for the varying representations of multiple stimuli that were reported in different regions of category-selective cortex in neuroimaging and single-unit recording studies. This type of organization may enable a dynamic and flexible representation of complex visual scenes that can be modulated by higher-level cognitive systems according to task demands.

## Introduction

A fundamental feature of primates’ high-level visual cortex is its division to category-selective areas, such as face, body or object-selective regions (Downing, Jiang, Shuman, & Kanwisher, 2001; Kanwisher, McDermott, & Chun, 1997; Kanwisher & Yovel, 2006; Malach et al., 1995; Peelen & Downing, 2005; Pinsk et al., 2009; Tsao, Moeller, & Freiwald, 2008). These category-selective regions reside in adjacent locations along the lateral occipital and ventral temporal cortex (Fisher & Freiwald, 2015; Grill-Spector & Weiner, 2014; Premereur, Taubert, Janssen, Vogels, & Vanduffel, 2016; Schwarzlose, Baker, & Kanwisher, 2005; Weiner & Grill-Spector, 2013). This division to areas selective to certain categories has led to numerous studies that have examined the profile of response of these category-selective areas to different categories when presented in isolation (e.g., Downing, Chan, Peelen, Dodds, & Kanwisher, 2006; Op de Beeck, Brants, Baeck, & Wagemans, 2010; Tsao, Freiwald, Tootell, & Livingstone, 2006; Yovel & Kanwisher, 2004). Nevertheless, visual scenes are typically composed of multiple objects. In the present study we demonstrate that this functional organization of neighboring clusters of category-selective neurons determines the representation of multi-category visual scenes.

The neural representation of multiple stimuli has been initially examined in single unit recording studies in low-level visual cortex. These studies have shown that the response to a preferred stimulus is reduced when presented with a non-preferred stimulus (e.g., Reynolds, Chelazzi, & Desimone, 1999, for review, Reynolds & Heeger, 2009). A normalization mechanism was proposed to account for these results. According to the normalization model, the response of a neuron to a stimulus is normalized by the response of its surrounding neurons to this stimulus (normalization pool) (Carandini & Heeger, 2012). When a preferred stimulus is presented together with a non-preferred stimulus, neighboring neurons that are selective to the non-preferred stimulus normalize the response of the neuron, resulting in a lower response to the pair of stimuli relative to the response to the preferred stimulus when presented alone.

Whereas the normalization model was initially developed based on the response of neurons in early visual cortex, findings supporting the normalization mechanism were also found in high-level visual cortex in both single-unit recording and fMRI studies. These studies examined the relative contribution of the isolated stimuli to the response of multiple stimuli and found different patterns of response in different areas of high-level visual cortex. The response of single neurons in inferotemporal cortex (IT) of the monkey (Zoccolan, Cox, & DiCarlo, 2005), as well as the fMRI response in object-selective cortex in humans (Baeck, Wagemans, & de Beeck, 2013; Macevoy & Epstein, 2009) to multiple stimuli, was the mean or a weighted mean response of the component stimuli. Unlike the mean response of general object areas, category-selective areas such as the face-selective or scene-selective areas in humans fMRI studies (Reddy, Kanwisher, & Vanrullen, 2009) as well as the face and body-selective neurons in monkeys (Bao & Tsao, 2018) showed a max response. In particular, the response to a multi-category stimulus composed of a preferred and non-preferred stimulus, was not reduced by the non-preferred stimulus but was similar to the response to the preferred category when presented alone (i.e. a max response). Bao & Tsao (2018) suggested that such pattern of response is consistent with the normalization model and can be explained by the degree of homogeneity of the normalization pool. If the surrounding neurons are selective to the same category as the recorded neuron (i.e., a face neuron in a face-selective area), the normalization pool is unresponsive to the non-preferred stimulus and therefore does not reduce the response of the recorded neuron to its preferred stimulus, yielding a max response. Taken together, previous single unit and neuroimaging studies have found either a mean response, a weighted mean response or a max response to multiple stimuli. These representations of multi-category stimuli may vary with the degree of homogeneity of the population of category-selective neurons (i.e. the homogeneity of the normalization pool) within a given cortical region and therefore reflect the operation of the same normalization mechanism in different areas of category-selective cortex (Bao & Tsao, 2018).

In the current study, we propose that category-selectivity, as measured with fMRI, can provide an estimate of the proportion of neurons that are selective to each of the measured categories and therefore with a measure of the homogeneity of the normalization pool. For example, a voxel that shows higher response to faces than bodies or objects has a larger proportion of face-selective than body or object-selective neurons (i.e. homogeneous normalization pool) (Tsao et al., 2006). A voxel that shows similar response to faces and bodies has roughly similar proportion of neurons that are selective to faces and bodies (i.e., heterogeneous normalization pool). Figure 1 shows the predictions of the normalization model for the response to a face and a body presented together in different cortical areas that are composed of face-selective neurons, body-selective neurons or with two populations of face-selective and body-selective neurons, as typically found in the borders between face- and body-selective areas. The response to multiple stimuli is expected to vary from a max response in areas with a homogeneous population of category-selective neurons to a mean response in an area with a mixed population of category-selective neurons (Fig. 1b). More generally, the normalization model predicts that the response to multiple stimuli is a weighted mean of the response to each of the stimuli, and that the weights are determined by the magnitude of category-selectivity to each of the stimuli (Fig. 1c). Thus, by using fMRI we can examine the variations in the representation of multi-category stimuli and their correspondence with the magnitude of category-selectivity across a large, continuous area of cortex.

**Figure 1:**
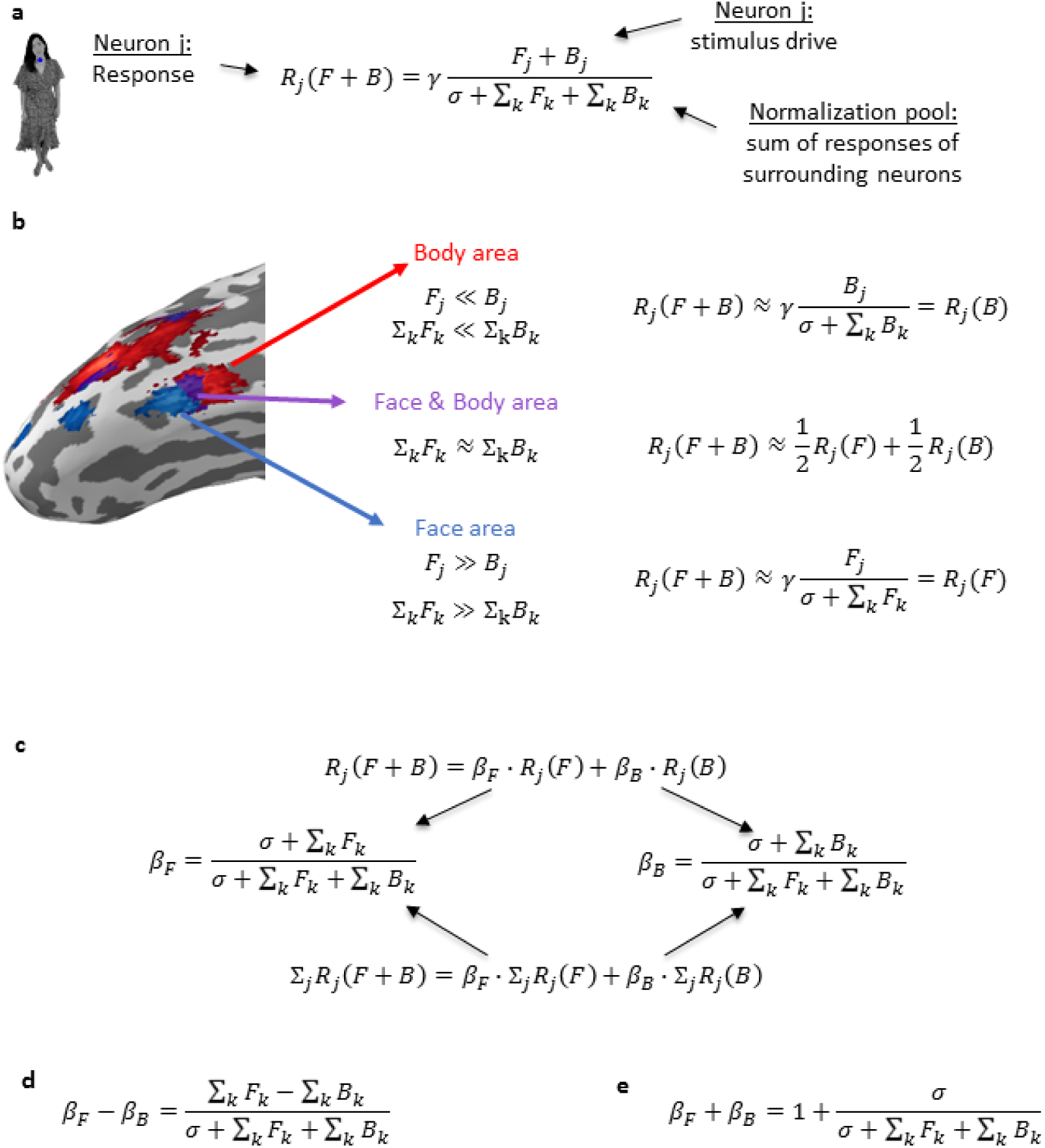
*(a) The normalization equation* (Reynolds & Heeger, 2009). *The response of a neuron is divided (normalized) by the sum of the responses of the surrounding neurons. Here we show the response to a face (F) and a body (B) presented together. (b) A surface map of face- and body-selective areas with the predicted response based on the normalization equation: a face-selective area (blue) and a body-selective area (red) contain homogeneous surrounding neurons that are selective to the same category, and therefore resulting in a max-like response. An area in the border between the face and body-selective areas (purple) contains a heterogeneous surrounding of face-selective neurons and body-selective neurons. If half of the neurons are face selective and half are body selective, then the response to a face and a body should be the mean of the responses to the isolated stimuli. (c) Using mathematical derivations of the normalization equation (a) (see Figure 1–figure supplement 1a for detailed derivation), the response to a pair of stimuli can be described as a weighted mean of the responses to the isolated stimuli. The weights (β*_*F*_ *and β*_*B*_*) are the contribution of the face and the body to the face+body response and are determined by the proportions of face and body-selective neurons within the normalization pool. The fMRI BOLD signal reflects the response of a sum of neurons with similar normalization pools, and therefore the same linear relationship between the pair and the isolated stimuli also applies for the fMRI response, with the same weights as for the single neuron equation (see Figure 1–figure supplement 1b). (d) The normalization equation further predicts that the difference between the weights corresponds to the difference in the proportions of face and body selective neurons, (e) and that the sum of weights is approximately 1 (see Figure 1–figure supplement 1c,d).* **Face image was replaced by illustrations due to bioRxiv’s policy on not including human faces within posted manuscripts.**

To test the correspondence between category-selectivity and the representation of multi-category stimuli in category-selective cortex, we ran two fMRI studies. In the first study we presented a face, a body and a face+body stimuli (Fig. 2a) and estimated the response to the combined stimulus based on the response to the isolated components by fitting a linear model to the data (Reddy et al., 2009). We found that category-selectivity to the isolated stimuli determines their contribution to the multi-category stimulus consistent with the predications stated above of the normalization model. In a second experiment, we replicated these findings and generalized them to a face+object stimulus (Fig. 2b).

**Figure 2:**
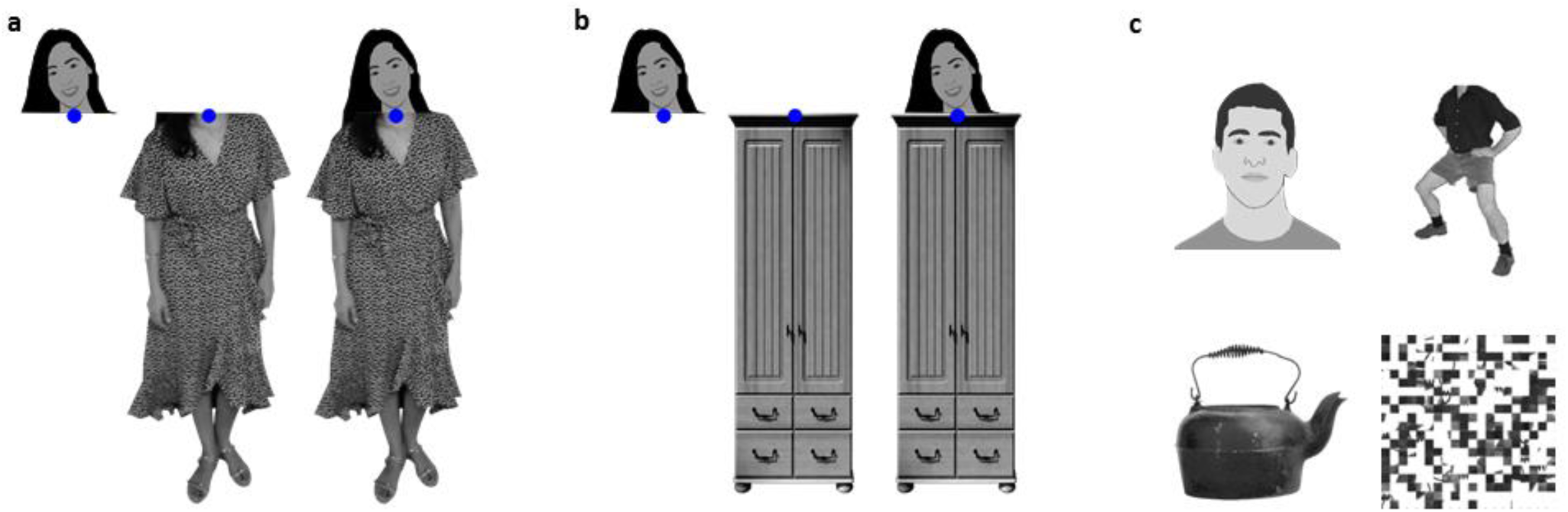
(a) A Face-Body stimulus set: face, body, and face+body stimuli, taken from the same images. **Face images were replaced by illustrations in this manuscript due to bioRxiv’s policy on not including human faces within posted manuscripts. The experiment stimuli included real human photos.** The fMRI response to these stimuli was used to estimate the contribution of the face and the body to the face+body representation. Participants were asked to fixate on the blue-dot and perform a one-back task (see Methods) (b) A Face-Object stimulus set: face, object, and face+object stimuli, all taken from the same images. Participants were asked to fixate on the blue-dot and perform a one-back task. We used wardrobes as the objects, which were matched to the body stimuli in terms of low-level visual properties. The fMRI response to these stimuli was used to estimate the contribution of the face and the object to the face+object representation. (c) Functional localizer stimulus set: faces, bodies, objects and scrambled objects. Functional localizer data were used to define category-selective regions of interest and to measure the voxel-wise selectivity to specific categories, independently from the data that were used to estimate the contribution of each part to the multi-category representation. The face and person images shown in the figures were not presented during the experiments but of individuals who gave consent to publish their images in this publication.

## Results

### Experiment 1 – The representation of a face+body in face- and body-selective areas

In an fMRI study, 15 subjects were presented with face, body and face+body stimuli, all taken from the same images (see Fig. 2a). They were instructed to fixate on the blue dot. In addition, these subjects were presented with a functional localizer, which included faces, bodies, objects (see Fig. 2c) and images of the whole person not used for the purpose of this study. The functional localizer data were used to define face and body-selective regions of interest (ROIs) as well as voxel-wise selectivity maps for faces and bodies. We estimated the contribution of the face and the body to the face+body response in these different category-selective areas by fitting a linear model to the data. Importantly, we did not limit the sum of the coefficients to 1 so they could take any value that best fits the data. Results support our predictions (Fig. 1) that category selectivity to the face and body in a given cortical area determines their contribution to the response to the combined face+body stimulus (Fig. 3). Moreover, we show that the entire cortical area that is selective to either faces or bodies follows the same principal normalization framework with the specific parameters determined by the local profile of category selectivity (Fig. 4).

**Figure 3:**
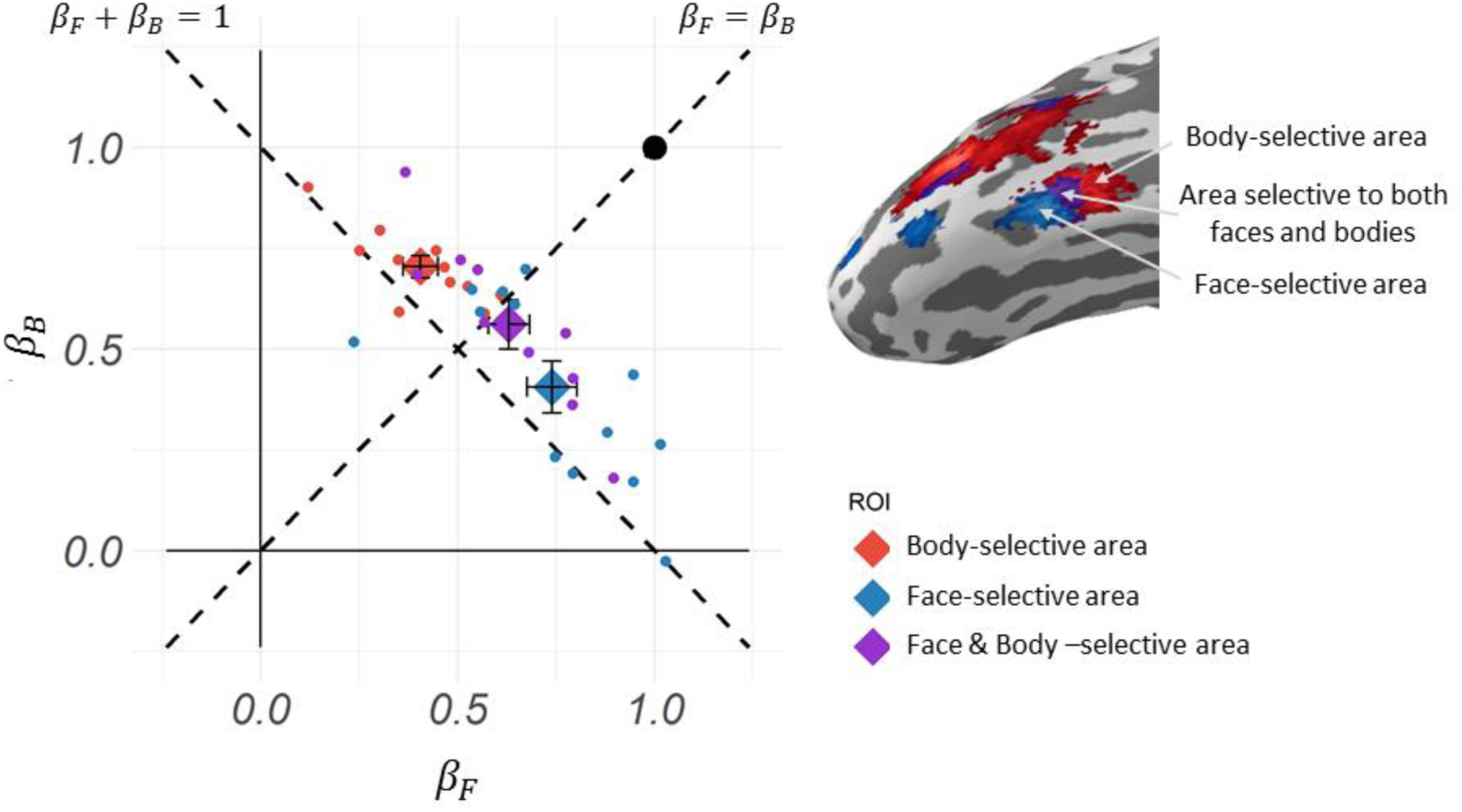
Experiment 1: Left: A scatterplot of the beta coefficients for the face and the body that best fit the response of the 30 most selective voxels within each subject’s ROI to the face+body stimulus. Each dot indicates the results of a single subject within an ROI (in the right hemisphere). β_F_ indicates the contribution of the face to the face+body response and β_B_ indicates the contribution of the body to the face+body response. The large diamonds indicate the group mean (error bars indicate s.e.m.). Right: a brain surface of one representative subject showing the location of the face-selective, body-selective and the overlap areas in lateral-ventral occipito-temporal cortex.

**Figure 4:**
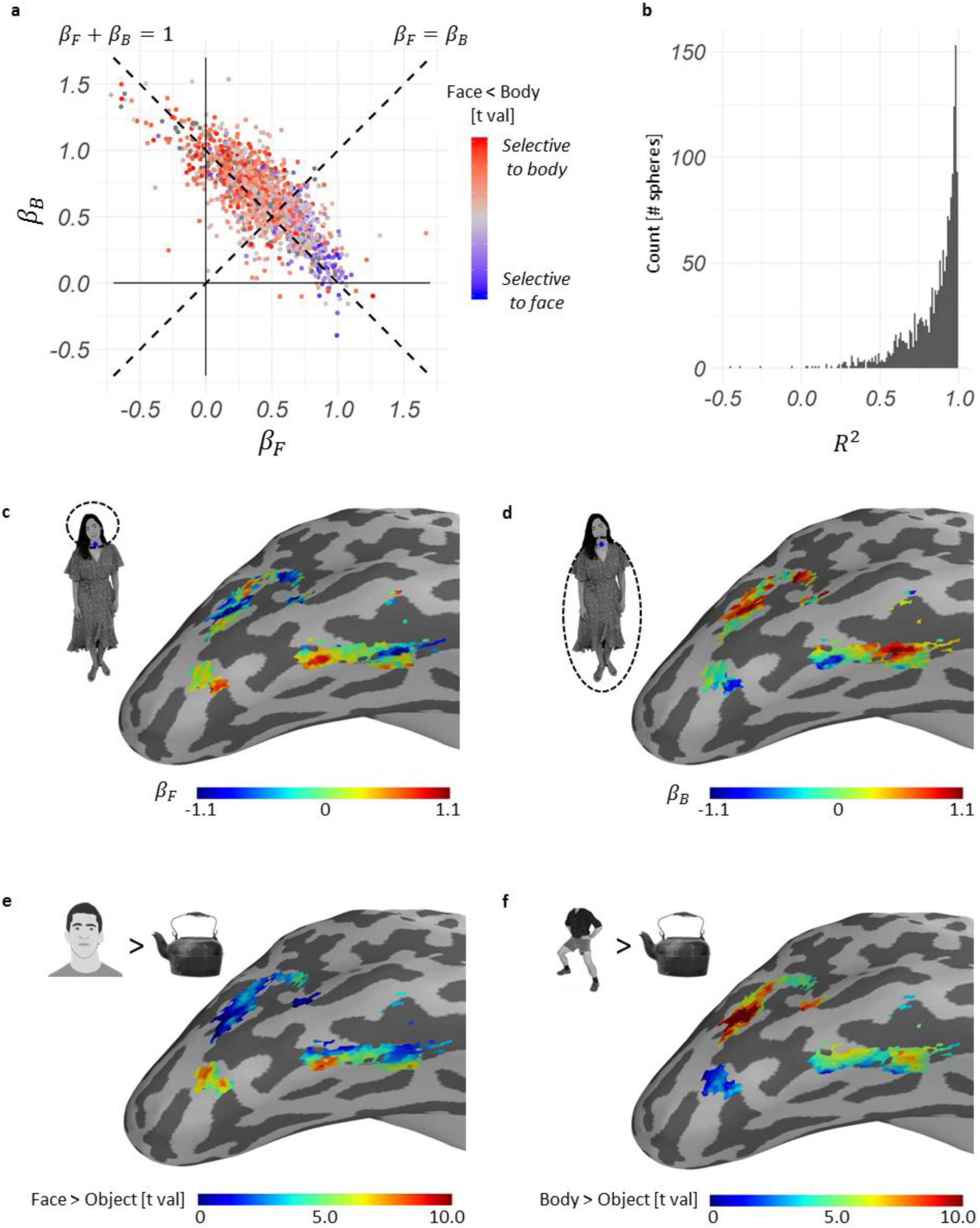
*Experiment 1: (a) The beta coefficients of all spheres of all subjects in the face and body selective areas indicating the contribution of the face (β*_*F*_*) and the body (β*_*B*_*) to the response to the face+body (equation (1)). The color of each dot indicates the selectivity for the face relative to the body based on independent functional localizer data. (b) A histogram of the R*^*2*^ *values of the linear models accounting for the response to the face+body of all spheres (negative values can be observed for models without intercept, see Methods) (c-f) Results of a representative subject plotted on the cortical surface for voxels that were selective to either faces or bodies: (c) The contribution of the face to the face+body representation as indicated by the face regression coefficients (β*_*F*_*). (d) The contribution of the body to the face+body representation as indicated by the body regression coefficients (β*_*B*_*). (e) Selectivity to faces (t map of Face>Object). Selectivity was determined based on independent functional localizer data. (f) Selectivity to bodies (t map of Body>Object). Selectivity was determined based on independent functional localizer data.* **Face images were replaced by illustration due to bioRxiv’s policy on not including human faces within posted manuscripts.**

#### Region of interest (ROI) analysis

First, we examined the contribution of the face and the body to the face+body response in the face and body-selective areas. For each individual subject, we extracted the face-selective area, body-selective area and the overlap between these areas (i.e. areas that are selective to both faces and bodies) (see Fig. 3 for an example of these areas in a representative subject). For each subject and each area within the right occipito-ventral cortex, we fitted a regression model for the response of the 30 most selective voxels (see Figure 3–figure supplement 1 for similar findings with different numbers of voxels) to predict the response to the face+body based on the responses to the face and the body (i.e., the percent signal change, PSC) in each of these voxels:

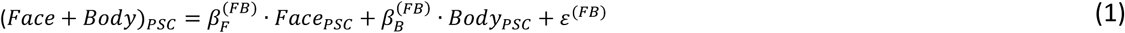

The beta coefficients 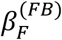 and 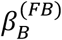 indicate the contribution of the face and the body to the face+body response for each area and each subject (The beta coefficients of the multi-category response model are not the same as the betas derived from the standard fMRI GLM analysis. The betas from the standard fMRI GLM analysis are used to determine the percent signal change (PSC) to each of the single- and multi-category stimuli as a measure of the fMRI response to that stimuli). All areas showed a significant contribution of both the face and the body to the face+body representation across all subjects, indicated by positive non-zero face and body coefficients (*β* = [0.39-0.74], p < .0001, see Fig. 2 and Figure 3–table supplement 1). Figure 3–figure supplement 2 shows similar results for the lateral occipital face and body areas, the Occipital Face Area (OFA) and Extrastriate Body Area (EBA).

Based on derivations of the normalization model we can further predict that the difference between the coefficients will correspond to the degree of selectivity of a cortical area to the different parts. In other words, the face coefficient should be higher than the body coefficient in face-selective areas, and vice versa for body-selective areas. (Fig. 1d. See Figure 1–figure supplement 1c for detailed derivation). Results were consistent with this prediction. We found that in the FFA, which is composed of mainly face-selective neurons, the contribution of the face was larger than the contribution of the body [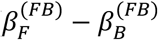: mean=0.334, t(12)=2.846, p=0.015]. Conversely, in the FBA, which is composed of mainly body-selective neurons, the contribution of the body was larger than the contribution of the face [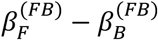: mean=-0.298, t(10)=-4.358, p=0.001]. In the area of overlap between the FFA and the FBA, which is selective to both faces and bodies, there was no significant difference between the contribution of the face and the body [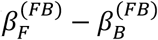: mean=0.070, t(10)=-0.628, p=0.544].

Additionally, we can further predict that the sum of the beta coefficients will be approximately 1 (Fig. 1e. See Figure 1–figure supplement 1d for detailed derivation). Indeed, the sum of weights was slightly over 1 [mean sum (s.e.m.): FFA: 1.145 (0.049); FBA: 1.110 (0.028); Overlap: 1.191 (0.024)] consistent with the normalization model predictions (Fig. 1e). In addition, the response to the face+body is consistent with a weighted mean response rather than an additive response, as indicated by the coefficients being smaller than 1 [all p-values <0.01], and the sum of these coefficients is lower than 2 [all p values <0.001]. Finally, we rule out an alternative explanation that the weighted mean response is due to saturation of the BOLD response to multiple stimuli. We found that 53.24% of the voxels in our data [FFA: 53.33%, FBA: 58.48%, Overlap: 47.88%.] showed higher response to one of the single stimuli (a face or a body) relative to the response to the combined stimulus (face+body).

To further assess if the weighted mean model (i.e., the normalization model, Fig. 1c) is the best fit to the data, we compared this model to two other models – one model containing a non-zero intercept and another model containing an interaction between the face and the body. We found that the model that best explains our results is a model with only the face and the body as predictors (see Table 1).

**Table 1:** Experiment 1 – Model comparison. In order to compare the proposed model predicted by the normalization equation (Fig. 1) to other models across all subjects, we used a Bayesian hierarchical model to predict the representation of the face+body stimulus based on the response to the face and the body. For each area we fitted three models (face and body; adding an intercept; adding an interaction). Values in the table indicate the Bayes Factor (BF) for the comparison between the model with only face and body factors to the other models, showing that this model best explain the results within all ROIs.

|  | Comparing models with and without intercept (BF) | Comparing models with and without interaction (BF) |
| --- | --- | --- |
| FFA | $2.14 \times 10^5$ | $1.94 \times 10^5$ |
| FBA | $3.45 \times 10^7$ | $5.36 \times 10^4$ |
| Overlap | 6.75 | $1.15 \times 10^4$ |

Next, we assessed the contribution of the face and the body to the face+body representation along the face and body areas within the right occipito-temporal and lateral areas (see Figure 4–figure supplement 1 for similar results of the left hemisphere). For each individual subject, we measured the response to face, body and the face+body stimuli of each voxel in these anatomical locations. We then applied a moving mask of a sphere of 27 voxels. For each sphere, we fitted a linear model to the responses of the voxels within the sphere to predict the response to the face+body based on the responses to the face and the body (Fig. 1c).

Figure 4a depicts the beta coefficients for the face and the body, i.e. the contribution of the face and the body to the face+body response, of all spheres within the face and body-selective cortices in the right occipito-temporal and lateral areas of all subjects. The coefficients are scattered along the weighted mean line, indicating a sum of coefficients of approximately 1 [mean sum=1.071, 95% confidence interval (C.I.): [1.036, 1.106], consistent with the normalization model prediction (Fig. 1e). Figure 4b displays the distribution of R^2^ of the models for all spheres indicating a good fit of the linear model to the data [median R^2^=0.90]. The color of each dot indicates the selectivity to the face relative to the body, as measured by the independent functional localizer. Furthermore, consistent with our predictions (Fig. 1d), the difference between the contribution of the face and the body to the face+body representation, (i.e. the difference between the beta coefficients) is correlated with the face and body-selectivity as measured by the independent functional localizer data. To examine the statistical significance of this correlation, the correlation was computed for each subject and transformed to a Z-Fisher score and the mean across subjects was compared to a null hypothesis of a correlation of zero [mean fisher z=0.458, t(14)=8.058, p<0.001, 95% C.I. (0.321, 0.595)]. To reduce statistical dependency in our dataset because of the overlapping moving mask, we used for the correlation analysis an interleaved mask, taking only spheres that their center is not immediately adjacent to another.

Figure 4c-d shows the same face and body coefficients presented in Fig. 4a of a single subject placed on a surface map of his brain. Figure 4e-f shows the distribution of category selectivity of the same subject within the same region for the face and the body as indicated by the independent functional localizer data (indicated in Fig. 4a by the color of the dots). Overall, Fig. 4 shows the correspondence between the selectivity and the contribution of the face and the body to the face+body representation throughout the continuum of the face- and body-selective regions: areas with high selectivity for faces and low selectivity for bodies show high contribution of the face to the face+body representation, while areas with low selectivity for faces and high selectivity for bodies show high contribution of the body to the face+body representation.

### Experiment 2 – The representation of a face+body and face+object in category-selective areas

In a second fMRI study, 15 subjects were presented with face, body and face+body stimuli, as well as face, object (wardrobe) and face+object stimuli forming a composite multi-category stimulus (see Fig. 2b). Similar to Experiment 1, we estimated the contribution of the face and the body to the face+body response as well as the contribution of the face and the object to the face+object response in different category-selective areas. In this study we first show a replication of findings of Experiment 1 with the face and body stimuli based on half of the data that was collected in Experiment 1 (3 runs instead of 6 runs). Then we show that the entire category-selective area follows the same normalization framework both for face+body and for face+object with the specific parameters determined by the local profile of category selectivity to the relevant categories.

For each individual subject we extracted the face-selective area, body-selective area and the object-selective area based on independent functional localizer data that presented faces, bodies, objects and scrambled objects. Each area was defined by voxels that show a significantly higher response to one category relative to the combined response of the three categories. Thus, in this ROI analysis, areas that are selective to two categories are excluded.

#### ROI analysis

First, we ran the same analysis reported above to examine the contribution of the face and the body to the face+body response in face- and body-selective areas. Findings replicated the results of Experiment 1 (Fig. 4A), with both the face and the body contributing to the response to the face+body [FFA: 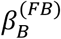: p=0.002. all other p-values <0.001, see Fig. 5a and Figure 5–table supplement 1]. Furthermore, the relative contribution of the face and the body varied as a function of the face and body selectivity (Equation 9), replicating the results of Experiment 1: in the FFA the contribution of the face was higher than the contribution of the body [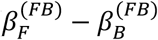: mean=0.494, t(14)=4.169, p<0.001], while in the FBA the contribution of the body was higher than the contribution of the face [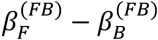: mean=-0.382, t(13)=-3.442, p=0.004]. The sum of coefficients in both face and body areas was again approximately 1 [mean sum (s.e.m.): FFA: 1.042 (0.066); FBA: 1.098 (0.054)] consistent with the normalization model predictions (Equation 10).

**Figure 5:**
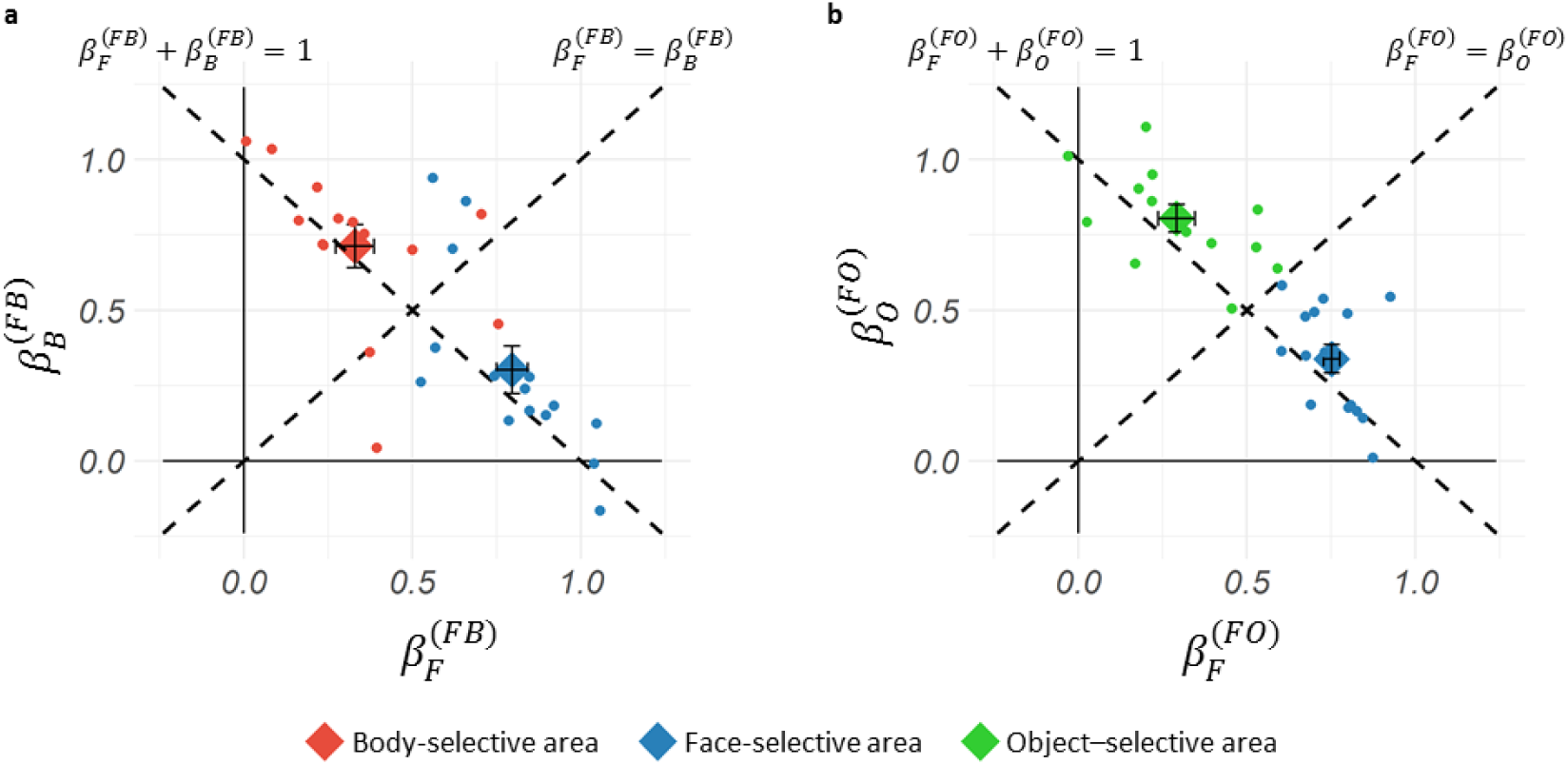
Experiment 2: (a) Beta coefficients for the face and the body predicting the response of the 30 most selective voxels within each subject’s ROIs to the face+body stimulus. 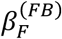 is the contribution of the face to the face+body response and 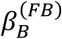 is the contribution of the body to the face+body response. Each dot indicates the results of a single subject within an ROI. The large diamonds indicate the group mean (error bars indicate s.e.m.). (b) Beta coefficients for the face and the object predicting the response of the 30 most selective voxels within each subject’s ROIs to the face+object stimulus. 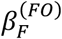 indicates the contribution of the face to the face+object response and 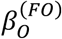 indicates the contribution of the object to the face+object response. Each dot indicates the results of a single subject within an ROI. The large diamonds indicate the group mean (error bars indicate s.e.m.).

Next, we performed similar analyses for the face+object stimuli. For each subject we fitted a regression model for the 30 most selective voxels within the face-selective area and the object-selective area to predict the response to the face+object based on the responses to the face and the object using the following equation:

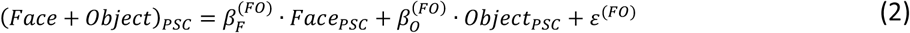

Similar to the face+body findings, the face- and object-selective areas showed a significant contribution of both the face and the object to the face+object representation across all subjects, indicated by positive non-zero coefficients of both the face and the object [all p-values<0.001, see Fig. 5b and Figure 5–table supplement 2]. In addition, the selectivity of the area determined the relative contribution of the face and the object to the face+object representation (Fig. 1d). Specifically, we found that in the FFA, which is mainly selective to faces, the contribution of the face was higher than the contribution of the object [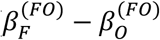: mean=0.413, t(14)=6.737, p<0.001], while in the object-selective area, the contribution of the object was higher than the contribution of the face [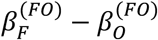: mean=-0.512, t(12)=-5.753, p<0.001]. The sum of coefficients, again, was 1 consistent with the normalization model (Fig. 1e) [mean sum (s.e.m.): FFA: 1.090 (0.043); Object area: 1.096 (0.047)].

The face+body stimuli are different from the face+object stimuli in that the former are a familiar combination whereas the latter are not. Previous studies have predicted different patterns of representations to familiar than non-familiar object combinations (Baldassano, Beck, & Fei-Fei, 2016; Kaiser & Peelen, 2018; Song, Luo, Li, Xu, & Liu, 2013) whereas others did not find such difference (Baeck et al., 2013; Kaiser, Strnad, Seidl, Kastner, & Peelen, 2014). To examine whether the pattern of response to face+body and face+object is different, we ran a repeated measure ANOVA with Pair Type (face+body, face+object) and ROI (face-selective, body/object selective) as within-subject factors and the difference between the coefficients as a dependent variable. We excluded from this analysis subjects that did not had 30 voxels in each of the three ROIs (3 subjects). As expected, the main effect of the ROI was significant [F(1,11)=54.382, p<0.0001], indicating that the selectivity of the ROI accounts for the relative contribution of each of the single categories to their multi-category stimuli. Importantly, we found no support for differences between Pair Type F(1,11)= 1.361, p=0.268, as well as no interaction between the ROI and Pair Type F(1,11)=0.024, p=0.808]. Thus, the same normalization framework accounts for the two types of multi-category stimuli.

#### Searchlight analysis

A similar searchlight analysis as described in Experiment 1 was performed for the face+body (equation (1)) and the face+object stimuli (equation (2)) in occipitotemporal and lateral areas that are selective to faces, bodies or objects relative to scrambled objects. Figure 5A depicts the beta coefficients for the face and the body, i.e. the contribution of the face and the body to the face+body response of all spheres within the category-selective cortices of all subjects. Although this area contains also voxels that are selective to objects, results are similar to Experiment 1. Specifically, the sum of coefficients is not significantly different from 1 [mean sum=1.013, t(14)=0.638, p=0.534, 95% C.I.=(0.970, 1.056)], and the difference in the contribution of the face and the body to the face+body representation, (i.e. the difference between the beta coefficients) is correlated with the selectivity to the face relative to the body as predicted [mean fisher z=0.407, t(14)=8.444, p<0.001, 95% C.I.=(0.304, 0.511)], replicating the results of Experiment 1.

We performed the same analysis for the face+object model over the same searchlight area and found similar results to the face+body findings (Fig. 6b): The beta-coefficients are scattered along the weighted mean line with a sum of coefficients that is not significantly different from 1 [mean sum=1.015, t(14)=1.490, p=0.158, 95% C.I.=(0.993, 1.038)] and the difference in the contribution of the face and the object to the face+object representation (i.e., the difference between the coefficients) is correlated with the selectivity to the face relative to the object as expected [mean fisher z=0.418, t(14)=11.193, p<0.001, 95% C.I.=(0.338, 0.498)] (Fig. 1 d-e).

**Figure 6:**
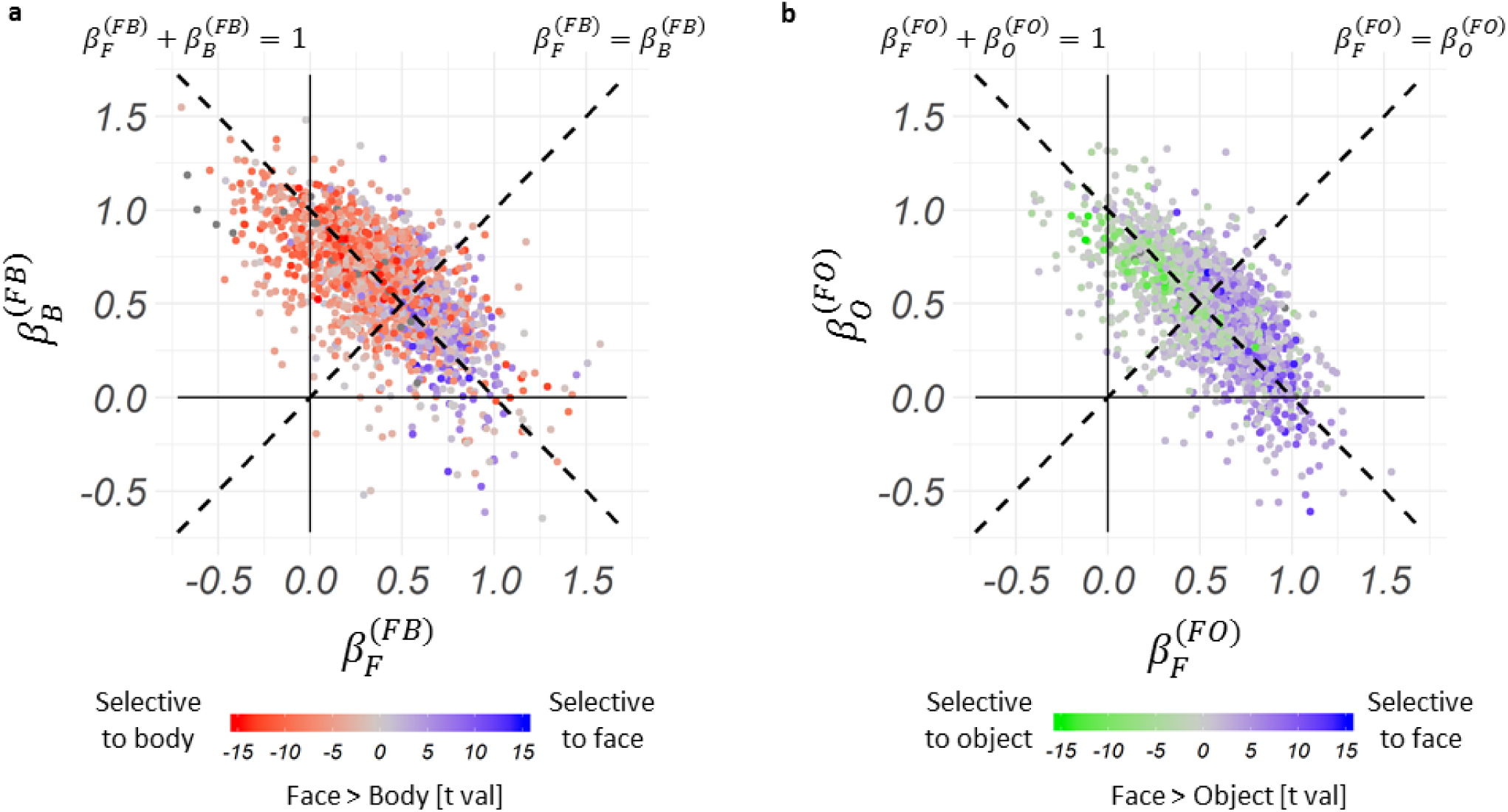
Results of searchlight analysis in Experiment 2. (a) The beta coefficients of all spheres in the category selective cortices of all subjects indicating the contribution of the face 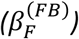 and the body 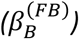 to the response to the face+body (equation (1)). The color of each dot indicates the selectivity for the face relative to the body based on independent functional localizer data. (b) The beta coefficients of all spheres in the category selective cortices (same as A) of all subjects indicating the contribution of the face 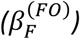 and the object 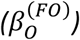 to the response to the face+object (equation (2)). The color of each dot indicates the selectivity for the face relative to the object based on independent functional localizer data.

To compare the spatial distribution of the beta-coefficients and category selectivity, we plotted the difference between the coefficients and the difference between the selectivity to each pair of categories on brain surface maps (Fig. 7a-d). Figure 7a shows the difference between the face and body coefficients (i.e., difference between the contribution of the face and the contribution of the body to the face+body representation) of one representative subject along his category-selective cortex. Figure 7b shows the selectivity to the face relative to the selectivity to the body for the same subject as measured by the independent functional localizer data. It can be seen that cortical areas that show higher contribution of the face to the face+body representation correspond to face-selective clusters (red in both figures), and that areas that show higher contribution of the body to the face+body representation correspond to body-selective clusters (blue in both figures). Figure 7c shows the difference between the contribution of the face and the object to the face+object representation for the same subject. Figure 7d shows the selectivity to the face relative to the object based on the functional localizer data. Similar to the face+body results, areas that show higher contribution of the face to the face+object representation correspond to face-selective clusters (red in both figures), and areas that show higher contribution of the object to the face+object representation correspond to object-selective clusters (blue in both figures).

**Figure 7:**
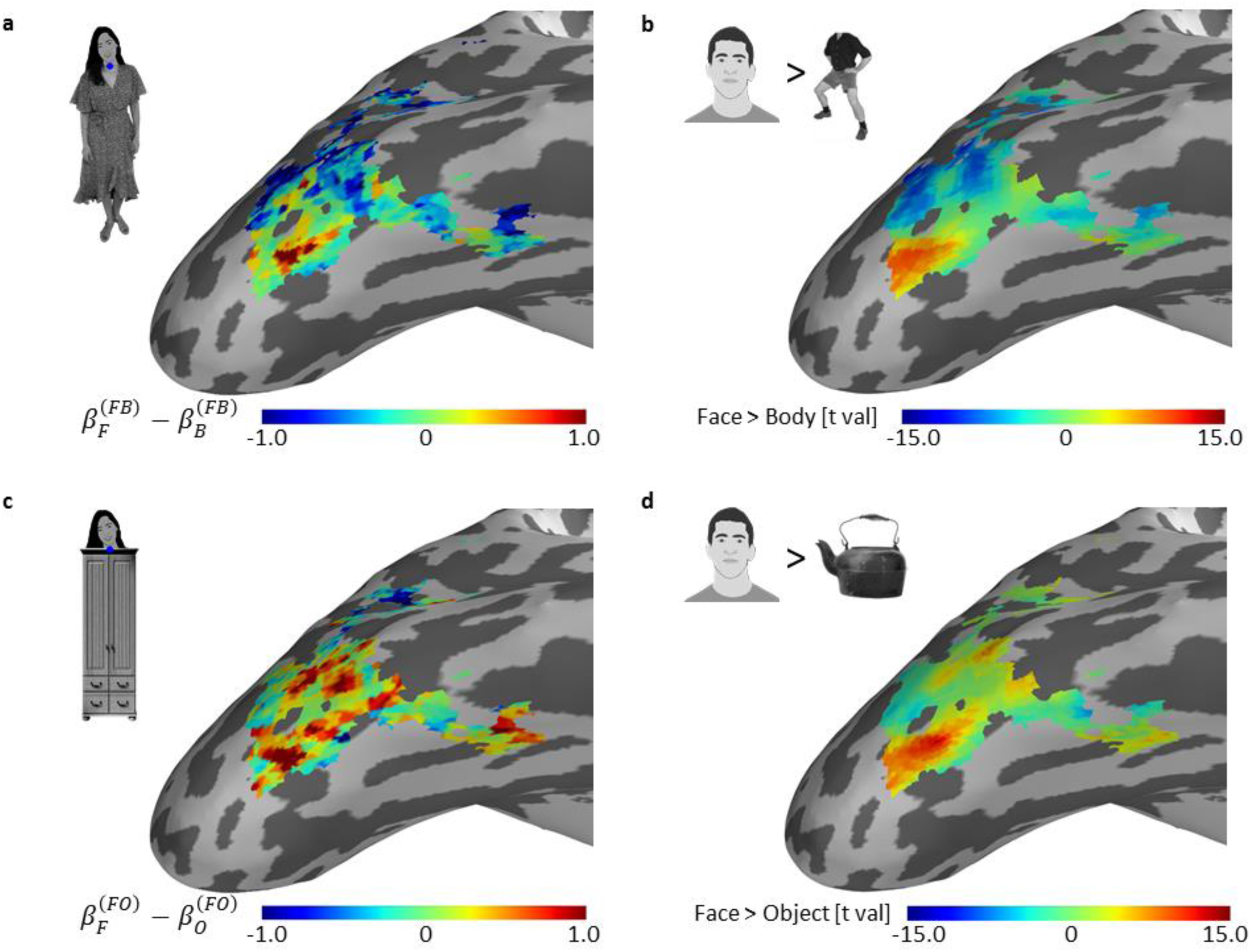
*Experiment 2: Results of searchlight analysis of one representative subject plotted on the cortical surface show the correspondence between the difference between the coefficients of the two categories and the magnitude of their selectivity. Note that Figure 3 shows a map of the coefficients and here we show a map of the difference between the coefficients. (a) The difference between the contribution of the face and the body to the face+body representation as indicated by the difference between the regression coefficients. A larger difference corresponds to a higher contribution of the face than the body to the representation of the face+body stimulus. (b) Selectivity to faces relative to bodies (t map of Face>Body). Selectivity was determined based on independent functional localizer data. (c) The difference between the contribution of the face and the object to the face+object representation as indicated by the difference between the regression coefficients. A larger difference corresponds to a higher contribution of the face than the object to the representation of the face+object stimulus. (d) Selectivity to faces relative to objects (t map of Face<Body) based on independent functional localizer data.* **Face images were replaced by illustrations due to bioRxiv’s policy on not including human faces within posted manuscripts.**

Finally, we computed the correlation between the beta coefficients and category selectivity for each category in category-selective cortex as well as a control area - early visual cortex. As expected, correlations between the beta coefficients and category-selectivity were found in category-selective areas but not in early visual cortex (see Fig. 8).

**Figure 8:**
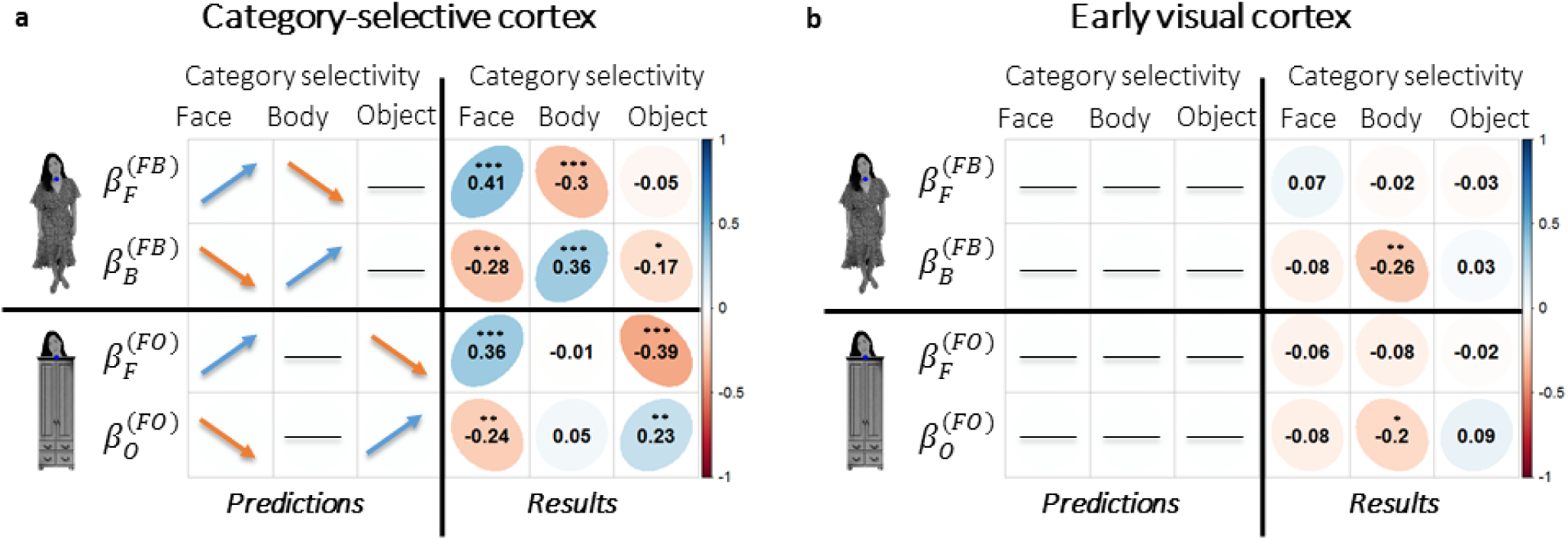
*Experiment 2 - Correlations between category-selectivity and model coefficients. Predictions and results of the correlations between the contributions of the single category to the multi-category response (beta coefficients of the models) and the category selectivity (based on the functional localizer data). (a) Results for category-selective cortex (Face, Body and Objects > Scrambled objects). A weighted mean model predicts that the contribution of each single category to the multi-category representation will be: (1) positively correlated to the selectivity to that same category; (2) negatively correlated to the selectivity to the other category that is present in the multi-category stimulus; and (3) not correlated with the selectivity to a category that is not present in the multi-category stimulus. Results are consistent with these predictions. (b) Results for early visual cortex (EVC). Since EVC does not show selectivity to object categories, we predict that category-selectivity will not be correlated with the contribution of the single object categories to the multi-category stimulus response. Values presented in the figure are the mean across subjects for the fisher z. *p<0.05, ** p<0.01, ***p<0.001 significant correlations corrected for multiple comparisons.* **Face images were replaced by illustrations due to bioRxiv’s policy on not including human faces within posted manuscripts.**

## Discussion

Our findings show that the functional organization of category-selective cortex determines the representation of multiple-category stimuli. Whereas previous studies have primarily focused on the role of this functional organization in the representation of single isolated stimuli (Grill-Spector & Weiner, 2014), visual scenes are typically composed of multiple stimuli. Thus, a great challenge of the visual system is to resolve competition among multiple stimuli (Cohen, Konkle, Rhee, Nakayama, & Alvarez, 2014; Kastner & Ungerleider, 2001; Peelen & Kastner, 2014) and generate a veridical representation of the objects that compose a complex visual scene. Here we show that by applying a normalization mechanism, the functional organization of neighboring clusters of category-selective neurons generates different representations along high-level visual cortex according to the profile of category selectivity. Areas with high concentration of neurons selective to a single category give priority to the preferred stimulus, filtering out the non-preferred stimuli. This operation enables hard-wired de-cluttering at early stages of visual processing (see also Bao & Tsao, 2018). Areas with a mixed population of category-selective neurons, enable similar, possibly competitive, representation to different categories that can be modulated by higher-level cognitive mechanisms according to task demands (Desimone & Duncan, 1995; Reynolds & Heeger, 2009).

The fMRI results reported in the current study are consistent with predictions derived from the normalization model that were developed based on single unit recording data (Fig. 1). The fMRI findings add to the neuronal findings by demonstrating the correspondence between the functional organization of high-level visual cortex and the representation of multi-category stimuli. This is enabled by two features of the fMRI signal: First, the magnitude of category-selectivity measured with fMRI provides a measure of the homogeneity of the normalization pool, an important factor in the normalization equation. Second, fMRI enables exploring the pattern of response across a large, continuous area of cortex. This pattern of response indicates that the representation of the multi-category stimulus changes gradually in a way that corresponds to the profile of category-selectivity (Fig. 4, 6, 7). These results propose a continuous mode of organization of high-level visual cortex, rather than the more common, discrete-like depiction of category-selective cortex.

Our findings provide a general framework that accounts for previous reports of single cell recording and neuroimaging studies that reported either a mean response (Macevoy & Epstein, 2009; Zoccolan et al., 2005), a weighted mean response (Baeck et al., 2013) or a max response (Bao & Tsao, 2018; Reddy et al., 2009) to multiple stimuli in different areas of category-selective cortex. We show that the relative contribution of each stimulus to the response of the compound stimulus varies along the weighted mean line and that this variation is accounted for by variation in category selectivity (Fig. 4, 6, 7). It is noteworthy that our findings **do not** imply that the representation of a multi-category stimulus of a single neuron is determined solely by *its own* selectivity to each of the stimulus categories (the nominator in the normalization equation). Category-selectivity measured with fMRI estimates the selectivity of the surrounding neurons (the denominator in the normalization equation) and therefore provides an estimate of the selectivity of the normalization pool and its effect on the response to multiple stimuli.

The normalization model was confirmed in our study by several measures: First, the model predicts a specific correspondence between the coefficients of the model and the selectivity of a cortical area, which was confirmed both in an ROI and a searchlight analysis. Second, we fit the data to alternative models, including a model with an interaction term and a model with a non-zero intercept, and found that the normalization model best accounts for the response to a multi-category stimulus (Table 1). Third, we performed the same analysis in early-visual cortex (Fig. 8), a control area that shows no selectivity to these object categories and therefore category-selectivity is not expected to explain the contribution of each of the single categories to the multi-category response, and indeed found no such relationship. Last, we ruled out an alternative explanation for the results, suggesting that the weighted mean is a result of the saturation of the BOLD signal to the multi-category stimuli, leaving the normalization model as the most probable explanation of our results.

Several neuroimaging studies that examined the representation of multiple stimuli have asked whether the response to a pair of stimuli deviates from a simple mean model, in particular for pairs of stimuli that show a meaningful relationship between them (Baldassano et al., 2016; Fisher & Freiwald, 2015; Kaiser & Peelen, 2017; Kaiser et al., 2014; MacEvoy & Epstein, 2011; Song et al., 2013). In these studies, a deviation from a simple mean response was considered as evidence for integration or a holistic representation of the complex stimulus. The main advantage of the linear model we used here is that it provides us with a direct measure of the type of deviation from the mean that the data show and can therefore decide between a weighted mean response, an additive response or a non-additive response. Our findings show that the deviation from mean reflects a weighted mean response. We found no evidence for a non-additive response to the combined stimulus and therefore no support for a holistic representation. This was the case both for the meaningful pair of face+body stimuli as well as for the non-meaningful face+wardrobe pair that generated similar representations. Similar results were reported by Baeck et al. (2013) that found the same representations for related and unrelated pairs of objects. Thus, the normalization mechanism operates in a similar manner for related and unrelated pairs of stimuli in object-category selective cortex.

Three additional studies that examined the representation of the whole person are noteworthy. Kaiser et al. (2014) reported no deviation from the mean in the response to a face and a body in a person-selective area (area defined by a whole person > objects). This area is likely to correspond to the overlap area reported in our study that is selective to both faces and bodies, and therefore consistent with our findings (Figure 2). Song et al. (2013) reported that only the right FFA showed a deviation from the mean for the response of the whole person and interpreted that as evidence for a holistic representation. This deviation, however, may reflect a weighted mean response rather than a non-additive response. Finally, Fisher & Freiwald (2015) examined the contribution of the face and body to the whole person in a monkey fMRI study and found a super-additive (more than the sum) response in anterior but not posterior face areas, in particular, in area AF in the dorsal bank of the superior temporal sulcus. The human analog of area AF is likely to be in the superior temporal sulcus (Yovel & Freiwald, 2013) an area that we did not examine in the current study that may apply a different mode of operation than the ventral visual cortex.

To summarize, our findings reveal a general framework of operation according to which the contribution of each stimulus to the representation of multiple stimuli in a given cortical area is determined by its profile of category selectivity, in line with a normalization mechanism. We therefore suggest that the functional organization of neighboring patches of neurons, each selective to a single or more categories, enables a flexible representation of complex visual scenes, where both de-cluttering and competition operate in different cortical areas, using the same type of neurons and the same mechanism of normalization. This type of organization may permit high-level cognitive processes to bias the response to any of these different representations according to task demands making the taxing operation of understanding complex visual scenes dynamic and flexible.

## Methods

### Participants

#### Experiment 1

Fifteen healthy volunteers (6 women, ages 19-37, 13 right-handed) with normal or corrected-to-normal vision participated in Experiment 1. Participants were paid $15/hr. All participants provided written informed consent to participate in the study, which was approved by the ethics committees of the Sheba Medical Center and Tel Aviv University, and performed in accordance with relevant guidelines and regulations.

The sample size (N=15) chosen for this study was similar to sample size of other fMRI studies that examined the representation of multiple objects in high-level visual cortex (10-15 subjects per experiment) (see for example: Baeck et al., 2013; Baldassano et al., 2016; Kaiser & Peelen, 2017; Kaiser et al., 2014; Macevoy & Epstein, 2009; MacEvoy & Epstein, 2011; Reddy et al., 2009; Song et al., 2013)

#### Experiment 2

Seventeen healthy volunteers (11 women, ages 20-30, 14 right-handed) that did not participate in Experiment 1, with normal or corrected-to-normal vision participated in Experiment 2. Two participants were excluded form analysis due to technical difficulties. Participants were paid $15/hr. All participants provided a written informed consent to participate in the study, which was approved by the ethics committees of the Sheba Medical Center and Tel Aviv University, and performed in accordance with relevant guidelines and regulations.

### Stimuli

#### Experiment 1

##### Main Experiment

Stimuli consisted of 40 grey-scale images of a whole person standing in a straight frontal posture with their background removed downloaded from the internet (20 men and 20 women identities). Each image of a person was cut into two parts approximately in the neck area resulting in a face stimulus and a headless body stimulus for each identity (Figure. 1A). The isolated face and body stimuli were presented in the same location they occupied in the whole person stimulus. A blue fixation dot was presented at a constant location around the neck on the screen across all conditions (at the center and upper part of the display) (Figure 1A). The size of the whole person image was approximately 3.5X12.2 degrees of visual angle.

##### Functional Localizer

Functional localizer stimuli were grey-scale images of faces, headless-bodies, non-leaving objects (Figure 1C), and images of the whole person that were not included in analyses of this study. The stimuli size was approximately 5.5×5.5 degrees of visual angle.

#### Experiment 2

##### Main Experiment

Experiment 2 contained two main parts: a face-body part and a face-object part. For the face-body part we used the same stimuli as in Experiment 1. For the face-object part we used pictures of faces, wardrobes and faces-above-wardrobes (Figure 1B). The face stimuli were the same 40 images of faces used in Experiment 1. For the object stimuli we used 40 images of grey-scale wardrobes with their background removed that were taken from the internet. We digitally manipulated the images of the wardrobes so that the object location, size (number of pixels on the screen), contrast and luminance will be matched to the 40 pictures of headless bodies from Experiment 1. The face+object stimuli were created by placing the wardrobe images right below the face in the same location of the body, i.e. a face above a wardrobe with no gap between them. A blue fixation dot was presented at a constant location on the screen across all conditions right over the neck in the same location as in Experiment 1) (Figure 1B). The size of the face+body stimuli as well as the face+object pair was approximately 3.5×12.2 degrees of visual angle.

##### Functional Localizer

Localizer stimuli were grey-scale pictures of faces, headless-bodies, non-leaving objects, and scrambled objects (Figure 1C). The size of the stimuli was approximately 5.5X5.5 degrees of visual angle.

### Apparatus and Procedure

#### fMRI acquisition parameters

fMRI data were acquired in a 3T Siemens MAGNETOM Prisma MRI scanner in Tel Aviv University, using a 64-channel head coil. Echo-planar volumes were acquired with the following parameters: repetition time (TR) = 2 s, echo time = 30 ms, flip angel = 82°, 64 slices per TR, multi-band acceleration factor = 2, acceleration factor PE = 2, slice thickness = 2 mm, field of view = 20 cm and 100 × 100 matrix, resulting in a voxel size of 2 × 2 × 2 mm. Stimuli were presented with Matlab (The MathWorks Inc.) and Psychtoolbox (Brainard, 1997; Kleiner et al., 2007) and displayed on a 32” high definition LCD screen (NordicNeuroLab) viewed by the participants at a distance of 155 cm through a mirror located in the scanner. Anatomical MPRAGE images were collected with 1 × 1 × 1 mm resolution, echo time = 2.88 ms, TR = 2.53 s.

#### Experiment 1

The study included a single recording session with six runs of the main experiment and three runs of functional localizer.

##### Main Experiment

Each of the six runs included 5 triads of face, body and face+body mini-blocks. Fig. 9 shows an example of two such triads. The order of face, body and face+body mini-blocks within each triad was counter-balanced across triads and runs. Each mini-block included eight stimuli of which 7 were of different identities and one identity repeated for the 1-back task. The identities presented in the face, body and face+body mini-blocks within a triad were different. Thus, each run included face, body and face+body stimuli of 35 different identities (7 identities x 5 triads). The 35 identities were randomly chosen from the set of 40 identities. Each mini-block lasted 6 seconds was followed by 12 seconds of fixation. A single stimulus display time was 0.325 s, inter-stimulus-interval was 0.425 s. Subjects performed a 1-back task (one repeated stimulus in each block). Each run began with a six seconds (3 TRs) fixation (dummy scan) and lasted a total of 276 seconds (138 TRs).

**Figure 9:**
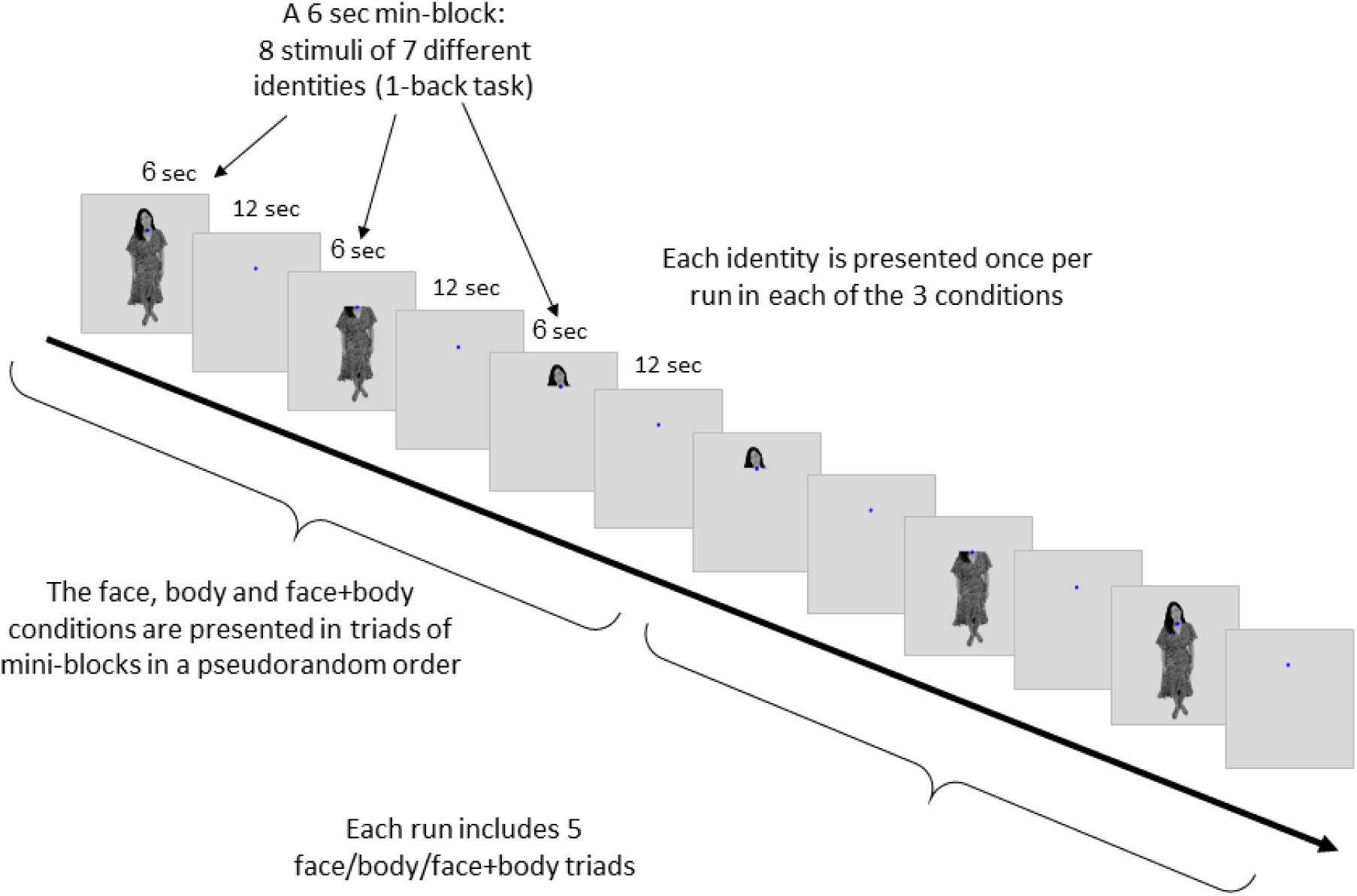
*Experimental procedure. Each run had 15 blocks of 3 conditions (5 blocks each). See Methods for a full description of the procedure***. Face images were replaced by illustrations in this manuscript due to bioRxiv’s policy on not including human faces within posted manuscripts. The experiment stimuli included real human photos.**

Subjects were instructed to maintain fixation throughout the run and their eye movements were recorded with an Eye tracker (EyeLink^®^).

##### Functional Localizer

Each run of the functional localizer included 21 blocks: 5 baseline fixation blocks and 4 blocks for each of the four experimental conditions: faces, bodies, objects and persons (analysis of person condition is not included in this paper). Each block presented 20 stimuli of 18 different images of which two repeated twice for a 1-back task. Each stimulus was presented for 0.4 sec with 0.4 sec Inter-stimulus interval. Each block lasted 16 seconds. Each run began with a six seconds fixation (3 TRs) and lasted a total of 342 seconds (171 TRs).

#### Experiment 2

The experiment included a single recording session with six runs of the main experiment and three runs of localizer.

##### Main experiment

The main experiment included 3 runs of face, body and face+body stimuli identical to Experiment 1. In addition, 3 runs of face, object and face+object stimuli were presented using the same design used for the face and body runs (Fig. 9). The face+object runs were presented before the face+body runs to avoid the priming of a body in the object and face+object mini-blocks. Subjects were instructed to maintain fixation throughout the run and their eye movements were recorded with an Eye tracker (EyeLink^®^). (See Figure 5 - Supplementary Figure 1 for the comparison of eye tracker data between conditions).

##### Functional Localizer

The functional localizer included four conditions of faces, bodies, objects and scrambled objects. All other parameters of the functional localizer runs were identical to Experiment 1.

## Data analyses

### fMRI Data Analysis and preprocessing

fMRI analysis was performed using SPM12 software, Matlab (The MathWorks Inc.) and R (R Development Core Team, 2011) costumed scripts, STAN (Carpenter et al., 2017) for Bayesian model fitting and Freesurfer (Dale, Fischl, & Sereno, 1999), pysurfer (https://pysurfer.github.io) and Python (http://www.python.org) costumed scripts for the surface generation and presentation. The code that was used for data analyses is available at https://github.com/LibiKl/multiple_objects_fMRI_analysis. The first three volumes in each run were acquired during a blank screen display and were discarded from the analysis as “dummy scans”. The data were then preprocessed using realignment to the mean of the functional volumes and co-registeration to the anatomical image (rigid body transformation). Spatial smoothing was performed for the localizer data only (5 mm). A GLM was run with separate regressors for each run and for each condition, including 24 nuisance motion regressors for each run (6 rigid body motion transformation, 6 motion derivatives, 6 square of motion and 6 derivatives of square of motion), and a baseline regressor for each run. In addition, a “scrubbing” method (Power, Barnes, Snyder, Schlaggar, & Petersen, 2012) was applied for every volume with frame-displacement (FD) > 0.9 by adding a nuisance regressor with a value of 1 for that specific volume and zeros for all other volumes. Percent signal change (PSC) for each voxel was calculated for each experimental condition in each run by dividing the beta weight for that regressor by the beta weight of the baseline for that run.

#### Experiment 1

##### Region of interest (ROI) analysis

Based on the functional localizer data, face- and body-selective voxels were defined individually for each subject using contrast *t*-maps. Regions of interest (ROI) were defined as clusters (>10 voxels) of voxels selective to a given category (p<10^−4^) within specific anatomical locations: (1) Fusiform face area (FFA): Face>Object within the Fusiform gyrus; (2) Fusiform body area (FBA): Body>Object within the Fusiform gyrus. The overlap areas were defined as the conjunction between face and body selective ROIs, included all voxels that were both face- and body-selective as described above. The 30 most selective voxels from each ROI within the right hemisphere were analyzed with the main experiment data. ROIs with less than 30 voxels were excluded from further analysis (see Figure 3–figure supplement 1 for the stability of the results across different number of voxels even with very low number of subjects).

##### Linear model fitting

The mean percent signal change (PSC) across runs to the face, the body and the face+body conditions from the main experiment data were extracted for each voxel within each ROI of each subject. For each subject, we fitted a linear model to predict the response to the face+body based on the response to the face and the body(Reddy et al., 2009) (equation 1). The features of the model were the response of a single voxel to a single condition. The sum of the coefficients was not pre-determined to sum up to 1 but was determined solely by fitting of the model to the data. We calculated the mean of the beta coefficients of the model, the mean difference between the coefficients and their mean sum across subjects.

To examine whether the linear model based on the normalization mechanism (Fig. 1c, equation 1) is the best fit to the data, we estimated a Bayesian hierarchical model to predict the response to a face+body based on the response to the face and the body including the data from all subjects for each ROI. In addition, we estimated two other Baysian hierarchical models: one with an addition of an intercept term, and another with the addition of an interaction between the face and the body. We then calculated Bayes factors to compare the models.

##### Univariate voxel-wise analysis

For each voxel within each ROI we compared the PSC to the face+body to the maximum PSC to the face and the body, and calculate the proportion of voxels that showed smaller response to the face+body, i.e., *face* + *body* < max(*face, body*). This analysis was done to assure that weighted mean response is not due to saturation of the BOLD response to face+body.

##### Searchlight analysis

For the searchlight analysis, we defined a face and body-selective region based on the localizer data by the contrast [(Face+Body)/2 > Object] (p<10^−4^) within the ventro-temporal and lateral occipital cortex. For each subject we defined a moving mask of a sphere of 27 voxels. To reduce statistical dependency in our dataset because of the overlapping moving mask, we used for the correlation analysis an interleaved mask, taking only spheres that their center is not immediately adjacent to another. For each sphere we fitted a linear model with its voxel data as features to predict the response to the face+body based on the response to the face and the body. The beta coefficients of these models represent the contribution of the face and the body to the response of the face+body of each sphere within the searchlight area. We then plotted a surface map of the beta coefficients of all spheres within the searchlight area to present the spatial distribution of the beta coefficients. We calculated R^2^ for each sphere and the median R^2^ across all spheres. Because the model we used does not include an intercept, negative R^2^ values indicate that a model is worse in predicting the dependent variable compared to a model that includes only an intercept.To examine the relationship between the difference between the face and body beta coefficients and the selectivity to face over a body (i.e., the t values of the contrast Face>Body from the independent functional localizer data) we performed a Pearson correlation across subjects. To assess the level of significance of the correlations, the correlation values were transformed to fisher’s Z, and a one-sample t-test was used against a null-hypothesis of zero.

#### Experiment 2

##### ROI Analysis

Based on the functional localizer data, face- body- and object-selective voxels were defined individually for each subject. Regions of interest (ROI) were defined as clusters (>10 voxels) of category selective voxels (p<10^−4^) within specific anatomical locations that show preference to a single category relative to all other categories: (1) Fusiform face area (FFA): Face > Body, Object & Scrambled-object within the Fusiform gyrus; (2) Fusiform body area (FBA):): Body > Face, Object & Scrambled-object within the Fusiform gyrus; (3) Ventral object area: Object > Face, Body & Scrambled-object within the medial part of the ventral temporal cortex. As in Experiment 1, the 30 most selective voxels from each ROI in the right hemisphere were chosen for model fitting. ROIs with less than 30 voxels were excluded from further ROI analysis.

The model fitting described in Experiment 1 was used to separately predict the response to the face+body based on the response to the face and the body (equation 1) and to predict the response to the face+object based on the response to the face and the object (equation 2). Similar to Experiment 1, we calculated the beta coefficients of the model, the mean difference between the coefficients and their mean sum for each model for each subject.

To examine whether the pattern of response to face+body and face+object is different, we ran a repeated measure ANOVA with Pair Type (face+body, face+object) and ROI (face-selective, body/object selective) as within-subject factors and the difference between the coefficients as a dependent variable. We excluded from this analysis subjects that did not had 30 voxels for all three ROIs (3 subjects excluded).

##### Searchlight analysis

For the searchlight analysis, we defined a category-selective region based on the localizer data by the contrast [(Face+Body+Object)/3 > Scrambled Object ((p<10^−4^)] within the Ventral-Temporal cortex and Lateral Occipital-Temporal cortex. A similar analysis that was performed in Experiment 1 was performed separately to the face and body runs and the face and object runs.

As a control area we also defined early visual cortex (EVC). EVC was extracted by performing an inverse normalization from an MNI space Brodmann area 17 mask to each subject’s native space. We matched the number of voxels in EVC to the number of voxels within the category-selective region by randomly choosing voxels from EVC. To further examine the correspondence between category selectivity and the contribution of each stimulus to the representation of the combined stimulus, correlations were computed between the selectivity to each of the three stimuli (face, body, object), measured by the t-value (each single category against all other categories), and the coefficients for each of the stimuli. The correlation values of all subjects were transformed to fisher’s Z, to examine the level of statistical significance against a null-hypothesis of a zero correlation. This analysis was performed in object-selective cortex and in early visual cortex (EVC).

## Data Availability

The code that was used for data analysis is available at https://github.com/LibiKl/multiple_objects_fMRI_analysis. Data that was collected in this study will be available after publication in a shared repository (https://openneuro.org/).

## Acknowledgments

This work is supported by a grant from the Israeli Science Foundation (ISF 446/16). We thank Tom Schonberg, Roy Mukamel, Jonathan Rosenblatt, Matan Mazor and Nathaniel Daw for helpful input on this work and Talia Brandman and Michal Bernstein for comments on this manuscript.

## Author Contributions

L.K. and G.Y. designed the experiments, interpreted the data, and wrote the paper. L.K. conducted the experiments and analyzed the data.

## List of supplementary material

### Supplementary figures

Supplementary Figure 1: Derivations of the normalization equation.

Figure 3–figure supplement 1: ROI analysis with different number of voxels.

Figure 3–figure supplement 2: Results for lateral face and body areas OFA and EBA.

Figure 4–figure supplement 1: Searchlight results for left hemisphere.

Figure 5-figure supplement 1: Eye tracker data distribution.

### Supplementary tables

Figure 3–table supplement 1: Experiment 1.

Figure 5–table supplement 1: Experiment 2 - Beta coefficients of face and body, ROI analysis.

Figure 5–table supplement 2: Experiment 2 - Beta coefficients of face and object, ROI analysis.

