## Supplementary Information for "The functional organization of high-level visual cortex determines the representation of complex visual stimuli"

**a**

$$\beta_F \cdot R_j(F) + \beta_B \cdot R_j(B) =$$

$$= \left(\overbrace{\frac{\cancel{\sigma + \Sigma_k F_k}}{\sigma + \Sigma_k F_k + \Sigma_k B_k}}^{\beta_F} \cdot \overbrace{\gamma \frac{F_j}{\cancel{\sigma + \Sigma_k F_k}}}^{R_j(F)}\right) + \left(\overbrace{\frac{\cancel{\sigma + \Sigma_k B_k}}{\sigma + \Sigma_k F_k + \Sigma_k B_k}}^{\beta_B} \cdot \overbrace{\gamma \frac{B_j}{\cancel{\sigma + \Sigma_k B_k}}}^{R_j(B)}\right) =$$

$$= \frac{\gamma F_j}{\sigma + \Sigma_k F_k + \Sigma_k B_k} + \frac{\gamma B_j}{\sigma + \Sigma_k F_k + \Sigma_k B_k} =$$

$$= \gamma \frac{F_j + B_j}{\sigma + \Sigma_k F_k + \Sigma_k B_k} = R_j(F + B)$$

$$\boxed{\Rightarrow R_j(F + B) = \beta_F \cdot R_j(F) + \beta_B \cdot R_j(B)}$$

**b**

$$\Sigma_j R_j(F + B) = \Sigma_j \gamma_j \frac{F_j + B_j}{\sigma_j + \Sigma_{k(j)} F_{k(j)} + \Sigma_{k(j)} B_{k(j)}} =$$

$$= \Sigma_j \left(\frac{\sigma_j + \Sigma_{k(j)} F_{k(j)}}{\sigma_j + \Sigma_{k(j)} F_{k(j)} + \Sigma_{k(j)} B_{k(j)}} \cdot \gamma_j \frac{F_j}{\sigma_j + \Sigma_{k(j)} F_{k(j)}}\right) + \Sigma_j \left(\frac{\sigma_j + \Sigma_{k(j)} B_{k(j)}}{\sigma_j + \Sigma_{k(j)} F_{k(j)} + \Sigma_{k(j)} B_{k(j)}} \cdot \gamma_j \frac{B_j}{\sigma_j + \Sigma_{k(j)} B_{k(j)}}\right) =$$

$$= \Sigma_j \left(\frac{\sigma_j + \Sigma_{k(j)} F_{k(j)}}{\sigma_j + \Sigma_{k(j)} F_{k(j)} + \Sigma_{k(j)} B_{k(j)}} \cdot R_j(F)\right) + \Sigma_j \left(\frac{\sigma_j + \Sigma_{k(j)} B_{k(j)}}{\sigma_j + \Sigma_{k(j)} F_{k(j)} + \Sigma_{k(j)} B_{k(j)}} \cdot R_j(B)\right) \approx$$

Under the assumption that the normalization pools of all neurons that are being summed are similar →

$$\approx \frac{\sigma + \Sigma_k F_k}{\sigma + \Sigma_k F_k + \Sigma_k B_k} \cdot \Sigma_j R_j(F) + \frac{\sigma + \Sigma_k B_k}{\sigma + \Sigma_k F_k + \Sigma_k B_k} \cdot \Sigma_j R_j(B) =$$

$$\boxed{= \beta_F \cdot \Sigma_j R_j(F) + \beta_B \cdot \Sigma_j R_j(B)}$$

**c**

$$\boxed{\beta_F - \beta_B} = \frac{\sigma + \Sigma_k F_k}{\sigma + \Sigma_k F_k + \Sigma_k B_k} - \frac{\sigma + \Sigma_k B_k}{\sigma + \Sigma_k F_k + \Sigma_k B_k} =$$

$$= \frac{\sigma + \Sigma_k F_k - \sigma - \Sigma_k B_k}{\sigma + \Sigma_k F_k + \Sigma_k B_k} =$$

$$\boxed{= \frac{\Sigma_k F_k - \Sigma_k B_k}{\sigma + \Sigma_k F_k + \Sigma_k B_k}}$$

**d**

$$\boxed{\beta_F + \beta_B} = \frac{\sigma + \Sigma_k F_k}{\sigma + \Sigma_k F_k + \Sigma_k B_k} + \frac{\sigma + \Sigma_k B_k}{\sigma + \Sigma_k F_k + \Sigma_k B_k} =$$

$$= \frac{\sigma + \Sigma_k F_k + \sigma + \Sigma_k B_k}{\sigma + \Sigma_k F_k + \Sigma_k B_k} =$$

$$= \frac{\sigma + \Sigma_k F_k + \Sigma_k B_k}{\sigma + \Sigma_k F_k + \Sigma_k B_k} + \frac{\sigma}{\sigma + \Sigma_k F_k + \Sigma_k B_k} =$$

$$\boxed{= 1 + \frac{\sigma}{\sigma + \Sigma_k F_k + \Sigma_k B_k}}$$

**Figure 1–figure supplement 1: Derivations of the normalization equation.** (a) The response to a face+body is represented by a linear combination of the responses to the isolated face and body. The beta coefficients are determined by the proportion of the neurons selective to either categories in the normalization pool. (b) The fMRI response to a face+body is estimated by the sum of the response of a group of neurons with similar normalization pools. (c) The difference between the beta coefficients is determined by the difference in the proportion of category-selective neurons in the normalization pool. (d) The sum of beta coefficients is approximately 1, suggesting a weighted mean model.

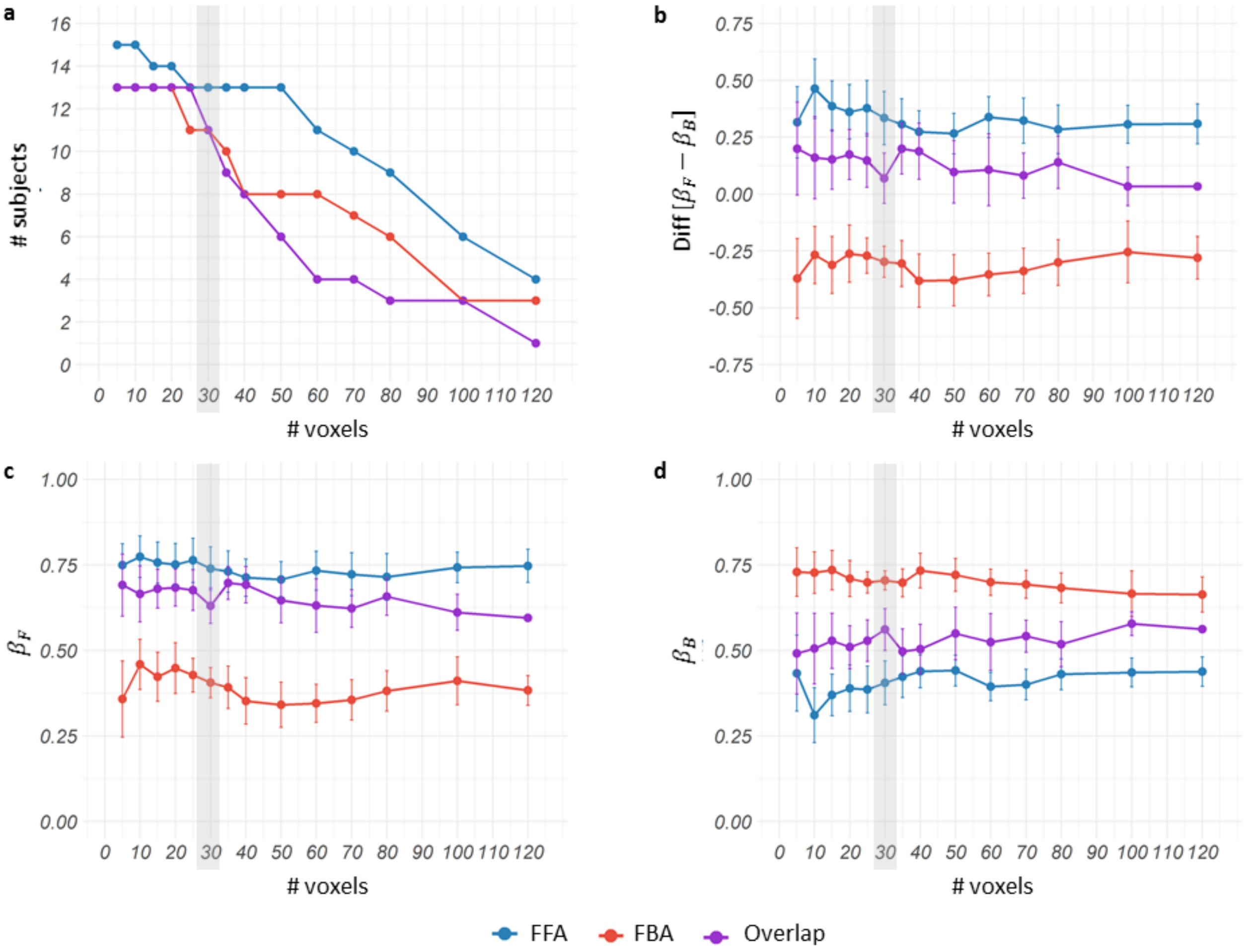


**Figure 3–figure supplement 1: ROI analysis with different number of voxels.** Analysis reported in the main text used 30 voxels for each ROI (marked in grey) (a) The number of subjects across different sizes of category-selective ROIs. As the size of the ROI grows the number of subjects decline. (b) The mean difference between the beta weights across subjects for each ROI size (error bars indicate s.e.m). (c) Mean $\beta_F$ and (d) mean $\beta_B$ across subjects for each ROI size. These data indicate that results are highly stable across different ROI sizes and number of subjects. Overall, results are highly stable across different number of voxels even when analysis includes very small sample sizes.

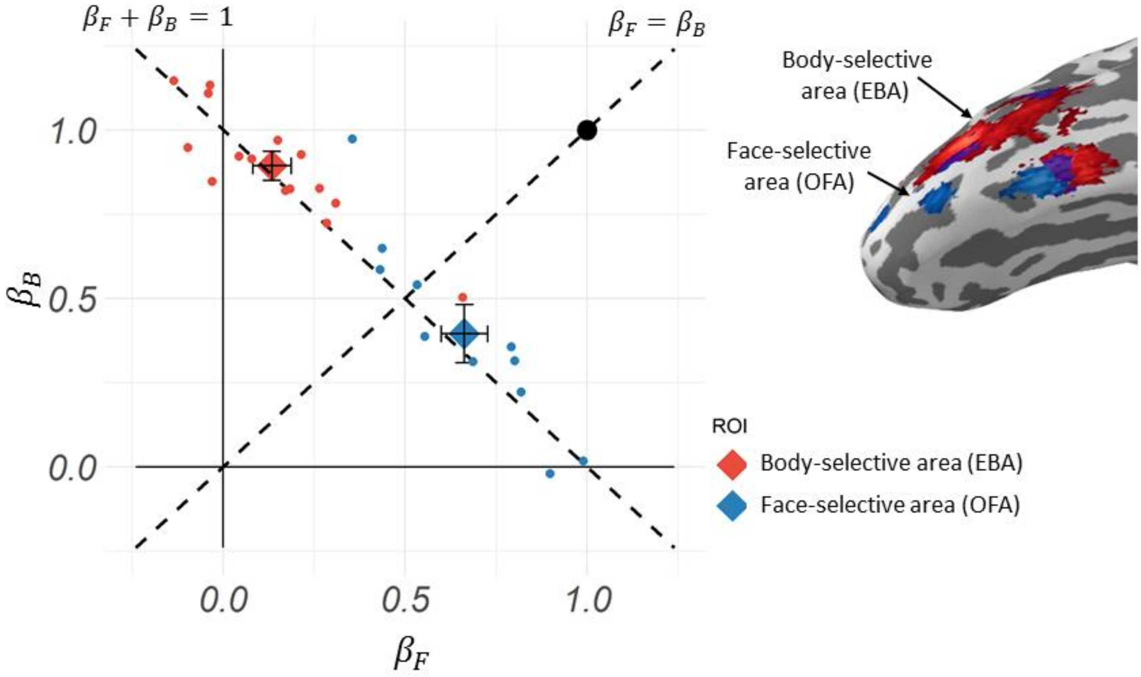


**Figure 3–figure supplement 2: Results for lateral face and body areas OFA and EBA.** Left: A scatterplot of the beta coefficients for the face and the body that best fit the response of the 30 most selective voxels within each subject's ROI to the face+body stimulus. Each dot indicates the results of a single subject within an ROI within the right hemisphere. $\beta_F$ indicates the contribution of the face to the face+body response and $\beta_B$ indicates the contribution of the body to the face+body response. The large diamonds indicate the group mean (error bars indicate s.e.m.). Right: a brain surface of one representative subject showing the location of the face-selective area (OFA), and body-selective area (EBA).

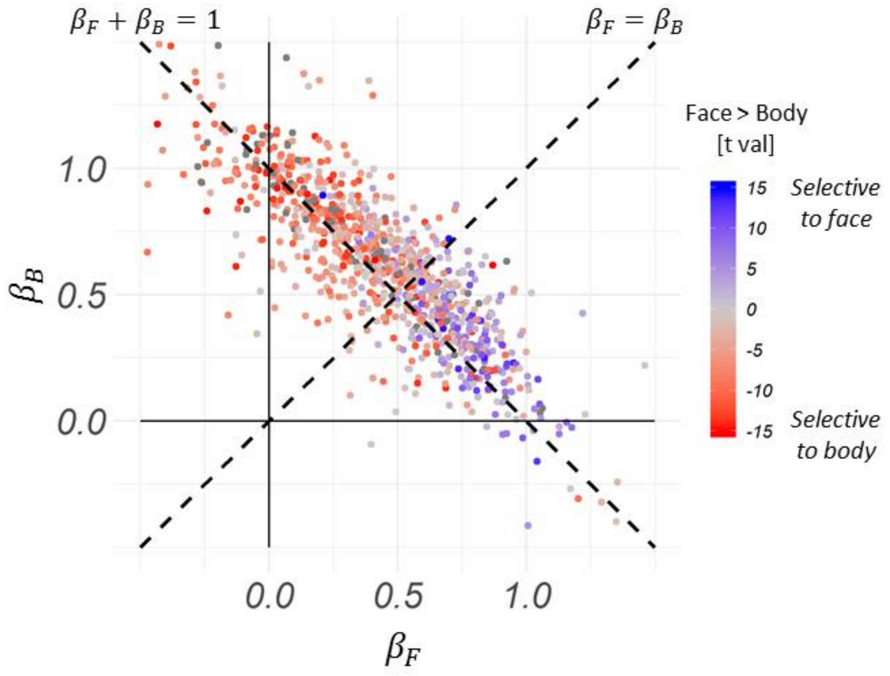


**Figure 4–figure supplement 1: Searchlight results for left hemisphere.** The beta coefficients of all spheres of all subjects in the face and body-selective areas indicating the contribution of the face ($\beta_F$) and the body ($\beta_B$) to the response to the face+body. The color of the dots indicate the selectivity of the face relative to the body, based on independent functional localizer data. Left hemisphere data show similar results to Right hemisphere data.

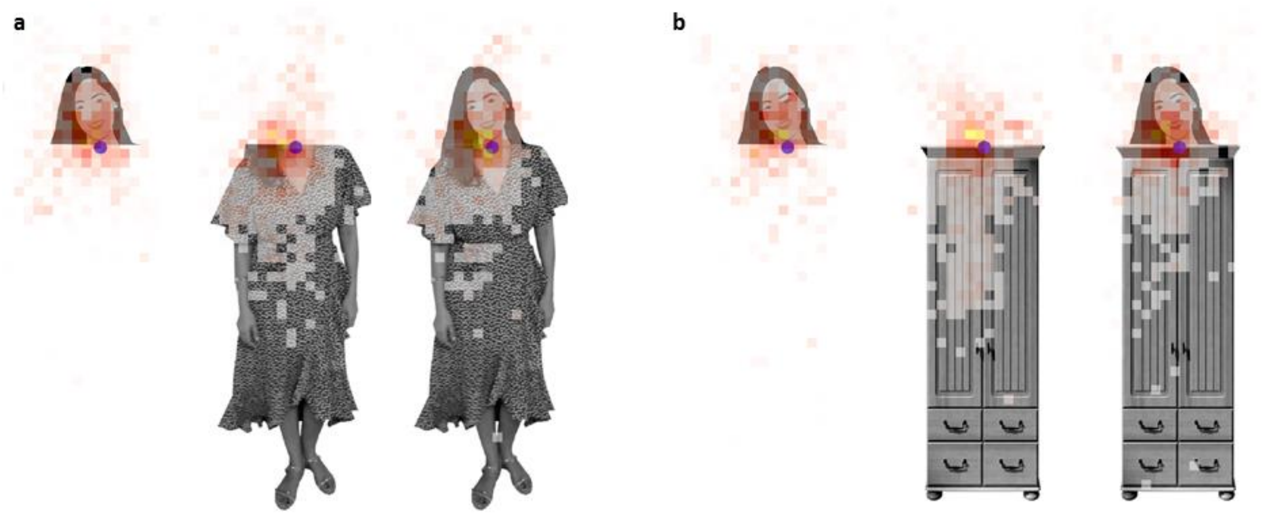


**Figure 5-figure supplement 1: Eye tracker data distribution.** The density distribution of fixations of all subjects are plotted over representative pictures of all experimental conditions. (a) Face+Body runs. (b) Face+Object runs. **Face images were replaced by illustrations in this manuscript due to bioRxiv's policy on not including human faces within posted manuscripts. The experiment stimuli included real human photos.**

| | Face coefficient - $\beta_F^{(FB)}$ mean (s.e.m.) | Body coefficient - $\beta_B^{(FB)}$ mean (s.e.m.) |
|---|---|---|
| **Face-selective area (n=13)** | 0.74*** (0.06) | 0.41*** (0.06) |
| **Body-selective area (n=11)** | 0.41*** (0.04) | 0.70*** (0.03) |
| **Face & Body selective area (n=11)** | 0.63*** (0.05) | 0.56*** (0.06) |

**Figure 3–table supplement 1: Experiment 1 - Beta coefficients of face and body, ROI analysis.** Mean and s.e.m. across subjects of beta coefficients for the face and the body the best fit the response of the 30 most selective voxels within each subject's ROIs to the face+body stimulus. *** p < .0001

| | Face coefficient - $\beta_F^{(FB)}$ mean (s.e.m.) | Body coefficient - $\beta_B^{(FB)}$ mean (s.e.m.) |
|---|---|---|
| **Face-selective area (n=15)** | 0.80*** (0.05) | 0.30* (0.08) |
| **Body-selective area (n=14)** | 0.33*** (0.06) | 0.71*** (0.07) |

**Figure 5–table supplement 1: Experiment 2 - Beta coefficients of face and body, ROI analysis.** Mean and s.e.m. across subjects of beta coefficients for the face and the body that best fit the response of the face+body based on the response of the 30 most selective voxels within each subject's ROIs. *p<0.1; ***p<0.0001.

| | Face coefficient - $\beta_F^{(FO)}$ mean (s.e.m.) | Body coefficient - $\beta_O^{(FO)}$ mean (s.e.m.) |
|---|---|---|
| **Face-selective area (n=15)** | 0.75*** (0.03) | 0.34*** (0.05) |
| **Object-selective area (n=13)** | 0.29** (0.05) | 0.80*** (0.05) |

**Figure 5–table supplement 2: Experiment 2 - Beta coefficients of face and object, ROI analysis.** Mean and s.e.m. across subjects of beta coefficients for the face and the object predicting the response of the face+object based on the response of the 30 most selective voxels within each subject's ROIs. **p<0.001; ***p<0.0001.
